# High-intensity physiological activation disrupts the neural signatures of conflict processing

**DOI:** 10.1101/2023.07.31.550835

**Authors:** Chiara Avancini, Luis F. Ciria, Clara Alameda, Ana Francisca Palenciano, Andrés Canales-Johnson, Tristan Bekinschtein, Daniel Sanabria

## Abstract

Physical activation fluctuates throughout the day. Previous studies have shown that during periods of reduced activation, cognitive control remains resilient due to neural compensatory mechanisms. In this study, we investigated the effects of high physical activation on both behavioural and neural markers of cognitive control. We hypothesized that while behavioural measures of cognitive control would remain intact during periods of high activation, there would be observable changes in neural correlates. In our electroencephalography study, we manipulated levels of physical activation through physical exercise. Although we observed no significant impact on behavioural measures of cognitive conflict, both univariate and multivariate time-frequency markers proved unreliable under conditions of high activation. Moreover, there was no modulation of whole-brain connectivity measures by physical activation. We suggest that this dissociation between behavioural and neural measures indicates that the human cognitive control system remains resilient even at high activation, possibly due to underlying neural compensatory mechanisms.

## INTRODUCTION

Fluctuations in physical and physiological activation occur naturally throughout the day. These spontaneous fluctuations unfold in a nonlinear manner ^1,2^ and become severe at extreme states such as deep sleep ^3^ or intense physical exertion ^4^, modulating cognitive processing ^5–8^. In particular, it has been shown that cognitive control (i.e., the ability to adaptively adjust cognitive processes to inhibit distracting information while maintaining task relevant information) is markedly impacted as the level of activation decreases ^3,9–11^. However, the majority of research on cognitive control has focused on fluctuations during decreased physical and physiological activation, with limited investigation into its effects under high physiological activation. In the current study, we aim at investigating the modulatory effect of high physical and physiological activation on behavioural and neural markers of cognitive control. To do so, we implemented an auditory conflict task combined with electroencephalography (EEG) recording in healthy human participants under a natural state of physical stress via exertion.

The interaction between heightened states such as physical exercise and cognitive performance in complex tasks does not appear to follow a linear relationship ^5^ and varies according to several variables such as the intensity of the exercise and the characteristics of the task. Some studies have reported better performance at medium intensity and worst at high intensity exercise ^12–15^. However, in the specifics of cognitive control tasks, physiological activation induced by exercise does not appear to hinder performance during acute exercise even if at high intensity ^12,16–18^. This is consistent with observations in the low end of the spectrum (drowsiness and sleep), which has received much more attention in the study of the interaction between physical/physiological activation and cognitive control. In fact, the ability to resolve cognitive conflict is not hampered by sleep deprivation and circadian rhythm ^9,10,19,20^, similarly to what observed during acute exercise. Interestingly, in the case of reduced activation, even if behavioural performance in conflict tasks appears to remain intact, univariate midfrontal (MF) theta and multivariate (congruency decoding) measures of cognitive conflict are no longer noticeable ^10^. Such discrepancy between behavioural and neural measures suggests that networks associated with the resolution of cognitive conflict may undergo reconfiguration when transitioning towards strained states. Does the neural reconfiguration observed during the resolution of cognitive conflict under decrease activation occur in conditions of high physical and physiological activation induced by exercise? In this pre-registered study ^21^, participants performed an auditory Simon task ^22^ during two bouts of aerobic exercise (cycling) at low and high intensity. We hypothesized that high-intensity exercise would not affect cognitive control at the behavioural level but it would disrupt univariate MF theta and multivariate spectral measures of cognitive conflict, similar to the case of drowsiness. In other words, we expected to gather evidence that under high intensity strained states the networks associated with conflict processing undergo reconfiguration. Consistent with our predictions, we found no behavioural difference in cognitive conflict measures between the two exercise conditions. However, neither the typical MF theta power signature of cognitive control nor the time-frequency multivariate decoding of conflict were reliable at high-intensity exercise. On the other hand, we found no difference between intensity levels in whole-brain connectivity measures. Therefore, we suggest that the human cognitive control system is resilient even at high physical activation states and propose that the dissociation between behavioural and neural measures could indicate the activation of neural compensatory mechanisms as a response to physiological pressure.

## RESULTS

### Bike and Physiological measures

To assess individual participants’ fitness, a maximum effort test was conducted on a stationary bike prior to the two EEG sessions. During the effort test, participants reached a mean maximum oxygen consumption (VO2_max_) of 3.374 l/min (SD = 0.478), a mean maximum heart rate (HR) of 180.589 bpm (SD = 13.224), and a mean maximum power output at VO2_max_ of 291.282 W (SD = 34.273).

Physiological measures were also obtained during the EEG sessions. During the low-intensity session, on average participants maintained 54.322% (SD = 6.422) of their maximum HR and moved 24.043% (SD = 6.109) of their maximum power output. During the high-intensity session, on average participants maintained 85.710% (SD = 4.225) of their maximum HR and moved 73.82% (SD = 6.18) of their maximum power output.

### Behavioural conflict and conflict adaptation effects

Behavioural responses to a Simon task were analysed to determine the effects of exercise intensity on cognitive conflict and conflict adaptation. Reaction times (RT) and accuracy rates (ACC) were recorded and analysed using repeated measure ANOVAs to compare performance across different intensity levels and trial conditions.

All participants showed positive conflict effects at both low and high-intensity (Figure 1A). To test the effect of physiological activation on the conflict effect we run a repeated measure ANOVA with intensity and congruency as factors. RT analyses revealed a main effect of congruency (F_(1,38)_ = 191.334, *p* < 0.001, 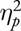 = 0.834, BF_10_ > 10000) with slower RT for incongruent than congruent trials (Figure 1B). Neither the main effect of intensity (F(1,38) = 0.554, *p* = 0.461, 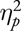 = 0.014, BF_10_ = 1.168) nor the interaction (F(1,38) = 2.304, *p* = 0.137, 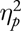 = 0.057, BF_10_ = 0.132) showed reliable differences. In ACC, the ANOVA revealed a main effect of congruency (F_(1,38)_ = 12.112, *p* < 0.001, 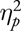 = 0.242, BF_10_ = 22.273) with higher accuracy to congruent than incongruent trials (Figure 1C). Neither the main effect of intensity (F(1,38) = 0.115, *p* = 0.736, 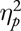 = 0.003, BF_10_ = 0.615) or the interaction (F(1,38) = 0.283, *p* = 0.598, 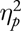 = 0.007, BF_10_ = 0.324) were statistically significant.

**Figure 1:**
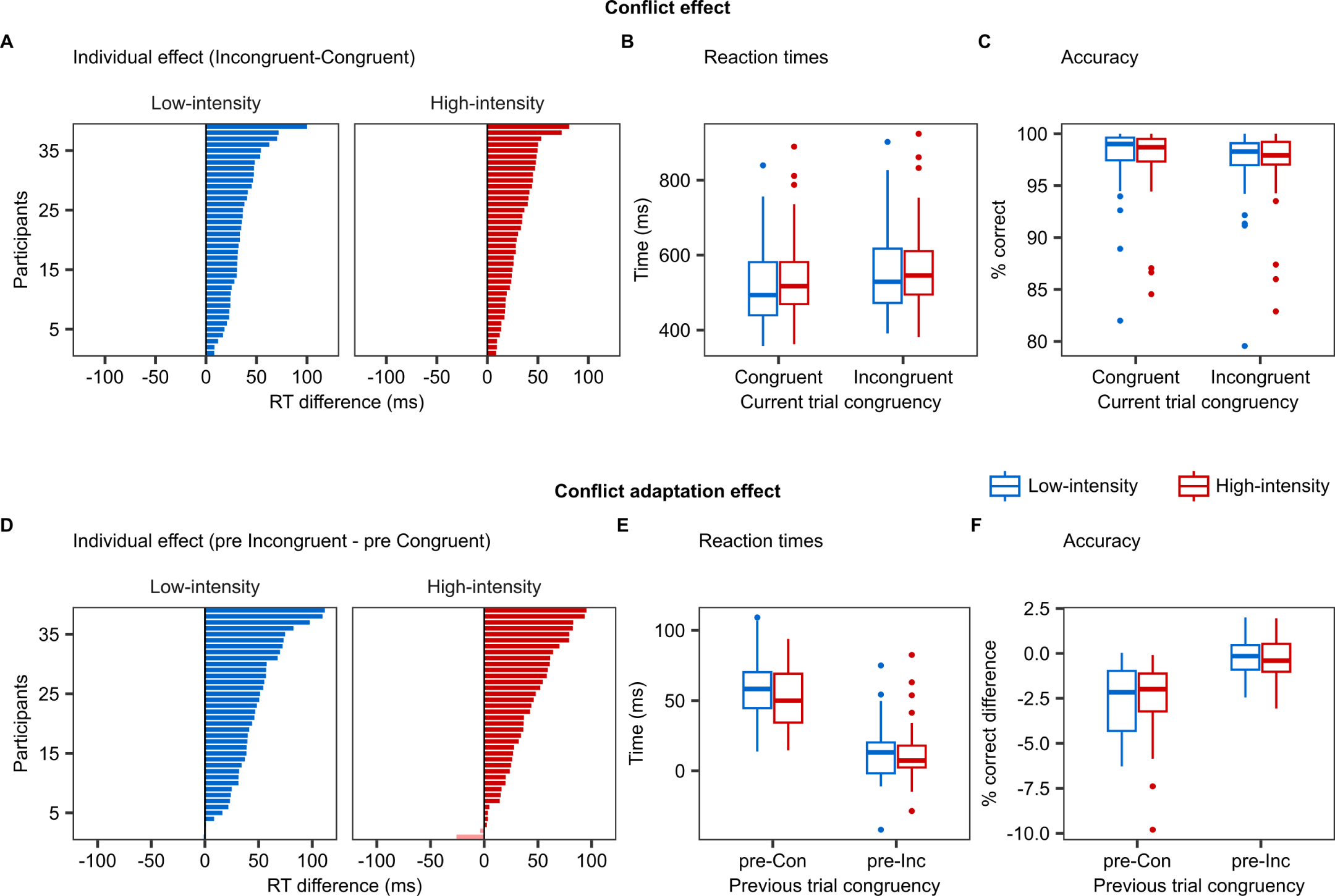
Behavioural results. **(A)** Individual conflict effect ordered according to increasing magnitude of the effect. **(B)** Boxplots illustrating the effect of intensity level on the reaction times (RT) of the responses to congruent (C) and incongruent (I) trials. Boxplots show the median (horizontal line within each box), the interquartile range (IQR; the box), and 1.5 times the IQR (whiskers). Outliers are indicated as individual points beyond the whiskers. **(C)** Boxplots illustrating the effect of intensity level on the accuracy (ACC) of the responses to congruent and incongruent trials. **(D)** Individual conflict adaptation effect ordered according to increasing magnitude of the effect. **(E)** Boxplots illustrating the effect of intensity level on the RT of the responses to trials preceded by congruent (pre-Con) and preceded by incongruent (pre-Inc) trials. **(F)** Boxplots illustrating the effect of intensity level on the ACC of the responses to trials preceded by congruent and preceded by incongruent trials.

The conflict adaptation effect was positive for 37 out of 39 participants at both low- and high-intensity (Figure 1D). To test the effect of intensity on the adaptation effect we run a repeated measure ANOVA with intensity and pre-congruency as factors and the behavioural conflict effect as dependent variable. In RT, there was a significant main effect of pre-congruency (F(1,38) = 134.735, *p* < 0.001, 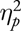 = 0.780, BF_10_ > 10000) with a smaller conflict effect when preceded by incongruent trials (Figure 1E). Neither the main effect of intensity (F(1,38) = 2.11, *p* = 0.154, 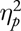 = 0.53, BF_10_ = 0.396) or the interaction (F(1,38) = 1.707, *p* = 0.199, 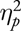 = 0.043, BF_10_ = 0.640) were statistically significant. In ACC, there was a main effect of pre-congruency (F(1,38) = 55.975, *p* < 0.001, 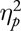 = 0.596, BF_10_ > 10000) with a smaller conflict effect when preceded by incongruent trials (Figure 1F). Neither the main effect of intensity (F(1,38) = 0.014, *p* = 0.908, 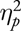 = 0.00036, BF_10_ < 0.0001) or the interaction (F(1,38) = 0.168, *p* = 0.684, 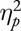 = 0.0040, BF_10_ = 0.250) were statistically significant.

### Univariate MF theta

Theta power changes were evaluated using time-frequency analysis of EEG data recorded during the cognitive task. The signal was decomposed between 2 and 30 Hz, and the time-frequency conflict effect was obtained by subtracting congruent from incongruent trial signals. Cluster-based permutation testing was used to examine interactions between congruency and intensity levels. The analysis on MF theta revealed significant difference between intensity conditions. This was shown by a positive cluster between 270-720 ms at 4-12Hz (Figure 2A). To explore what drove the effect we performed t-test against zero in low- and high-intensity on the power within the cluster area (Figure 2B; note that this additional analysis had not been preregistered). The one-sample t-test against 0 was significant in low-intensity (t(38) = 7.233, *p* < 0.001, Cohen’s d = 1.16, BF_10_ > 10000). On the other hand, a conflict effect was not reliably detected in high-intensity as shown by both frequentist and bayesian statistics (t(38) = 1.985, *p* = 0.054, Cohen’s d = 0.318, BF_10_ = 1.010). The power within the region of interest (ROI) showed positive conflict effect in 34 out of 39 participants at low-intensity and 26 out of 39 at high-intensity (Figure 2C). Therefore, as predicted, the difference in conflict effect between intensity levels was significant for the neural signature of MF theta, suggesting that physiological activation modulates the processing of cognitive conflict. Moreover, the effect was driven by a reliable conflict effect during the control condition of low-intensity but not during high-intensity. The repeated-measures ANOVA on average theta power within the interaction ROI showed a main effect of congruency (F(1,38) = 32.860, p = < 0.001, 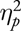 = 0.464, BF_10_ = 5.105) and an interaction between intensity and congruency (F(1,38) = 20.104, p < 0.001, 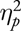 = 0.346, BF_10_ = 477.569), but no reliable main intensity effect (F(1,38) = 3.962, p = 0.054, 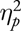 = 0.346, BF_10_ = 1.548). In pairwise comparisons, we found no significant difference between intensity levels in the congruent condition (t(38) = -0.400 p = 0.691, Cohen’s d = 0.064, BF_10_ = 0.189). On the other hand, we did find a statistically significant difference in intensity levels in the incongruent condition (t(38) = -3.18, p = 0.003, Cohen’s d = 0.509, BF_10_ = 11.861. Figure 2D). These results mirror the effects found in the drowsy state ^10^.

**Figure 2:**
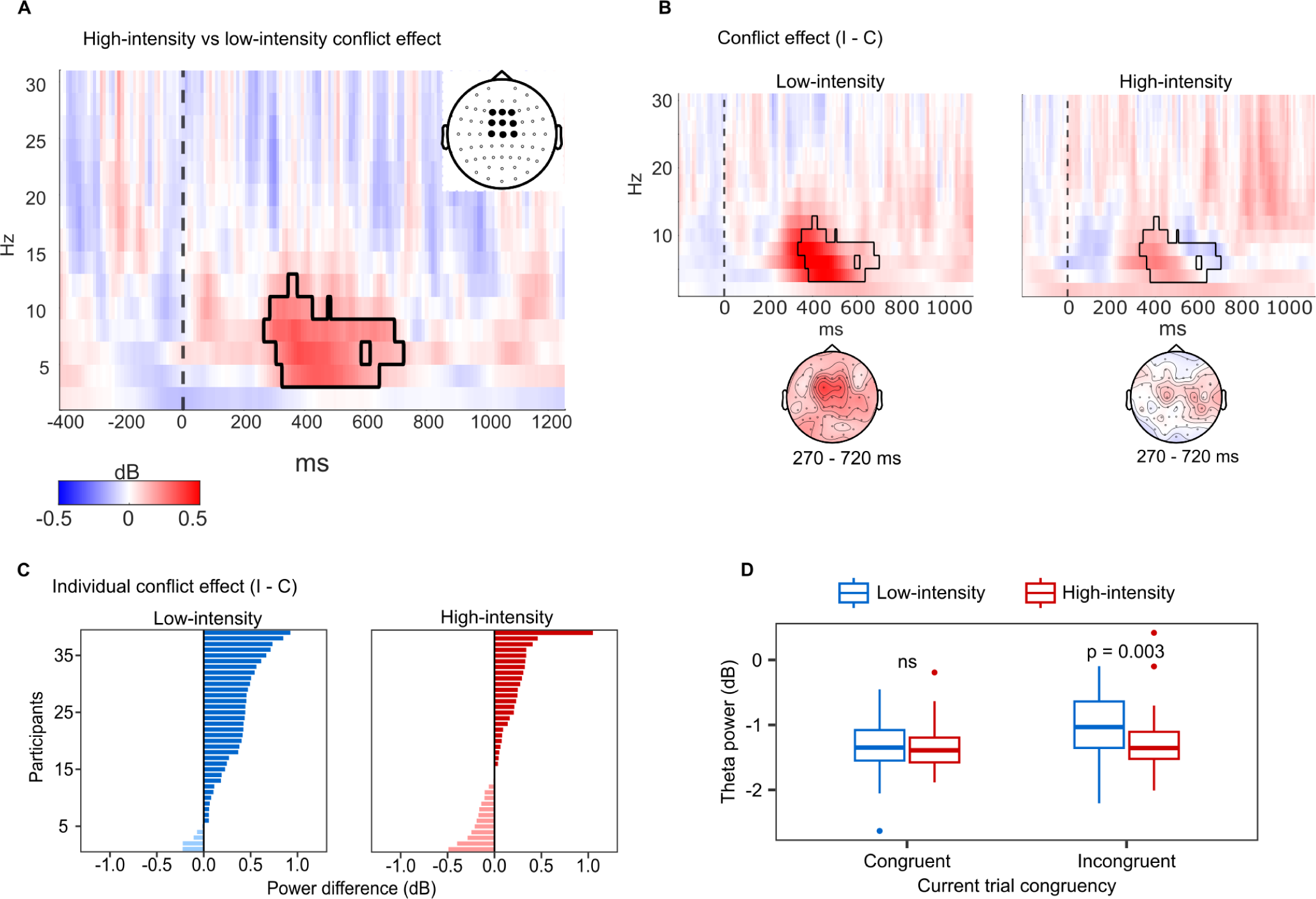
Univariate spectral analysis of mid-frontal theta-band oscillations. **(A)** Cluster permutation analysis capturing the difference between the conflict effect at high-intensity and the conflict effect at low-intensity. The solid black line delineates the significant positive cluster with *α* = 0.05. Bold dots in the top-right headplot represent the region of interest included in the analysis. **(B)** Conflict effect computed by subtracting the congruent (C) from the incongruent (I) condition separately for low- and high-intensity. The topographies represent theta-band power distribution within the timing of the significant cluster, which is delineated by a solid black line. **(C)** Individual mid-frontal (MF) theta conflict effect. For each participant, the power of the conflict effect was extracted from the area of the significant cluster as delineated in the interaction plot and then ordered according to increasing values. The lighter shade represent a negative conflict effect. **(D)** Boxplots illustrating the effect of intensity level on MF theta power of the responses to congruent and incongruent trials. Boxplots show the median (horizontal line within each box), the interquartile range (IQR; the box), and 1.5 times the IQR (whiskers). Outliers are indicated as individual points beyond the whiskers. The difference between intensity conditions was significant (p = 0.003) in incongruent trials but not significant (ns) in congruent trials.

We conducted exploratory analyses to investigate the relationship between behavioural measures and MF theta power changes induced by physiological activation (Figure 3). We fitted hierarchical linear mixed models with RT as dependent variable and intensity level and MF power changes as predictors. Exploratory analyses further support that at high-intensity MF theta does not bear relationship with behavioural measures of cognitive control as suggested by the lack of correlation between RT and MF theta power changes (Figure 3D). Indeed, the intercept was the best fitting model, having the lowest AIC (659.19) and BIC (666.26). Furthermore, we explored the possibility that MF power changes are predicted by latent automatic (task-irrelevant) processes underlying decision making. We used Drift Diffusion Modelling for conflict tasks on the behavioural data (DMC ^24^) to estimate the influence of automatic processes on decision time ^23^ and fitted hierarchical linear mixed with MF theta power as dependent variable and state and the automatic estimate as predictors.

**Figure 3:**
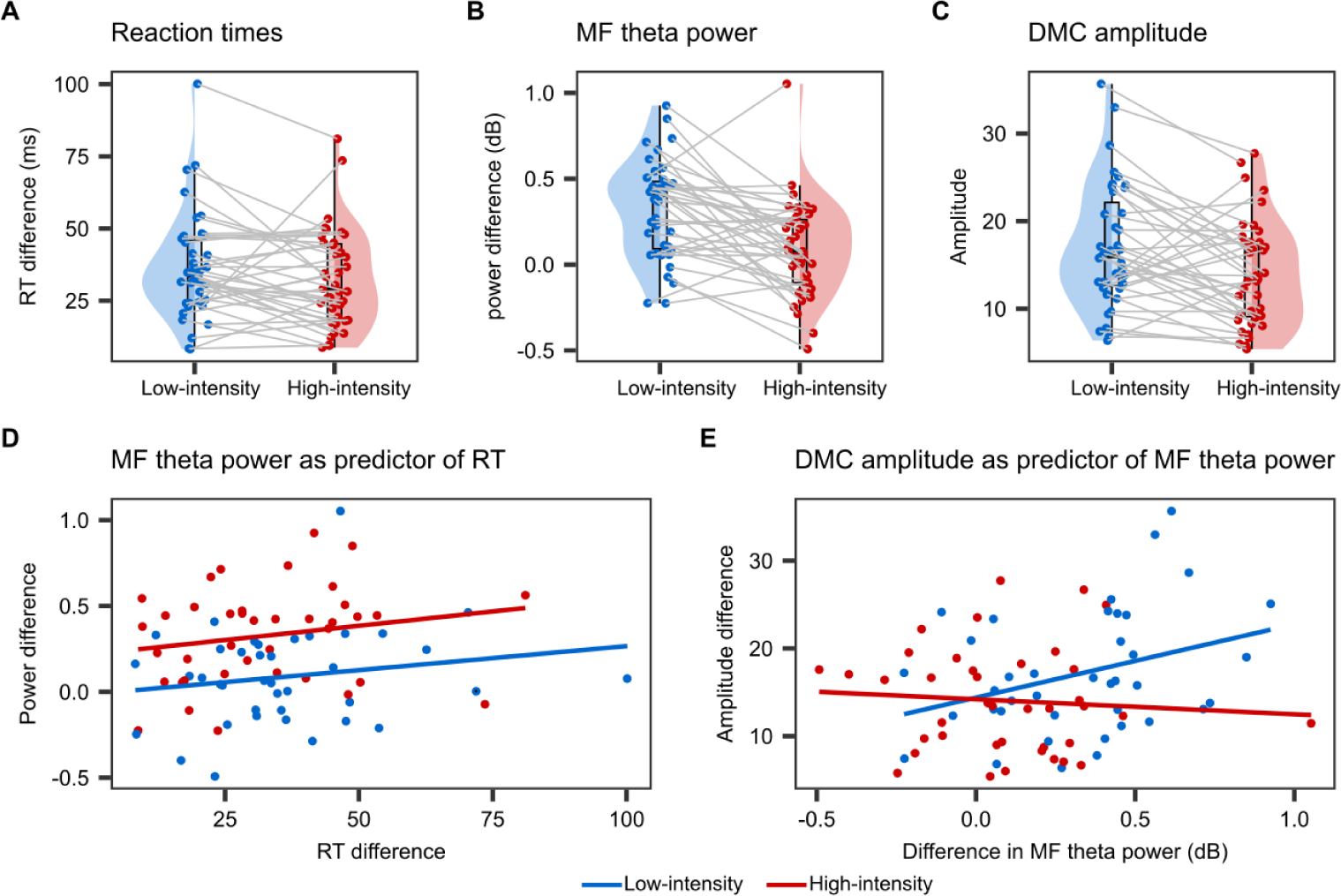
Visualization of individual differences across intensity levels and fitted linear models. Subject-by-subject differences elicited by intensity level manipulation. Blue and red areas are rotated kernel density plots. Box plots visualise the median (middle horizontal line), and 25th and 75th percentiles (bottom and top horizontal lines). The upper and lower whiskers indicate the distance of 1.5 times the interquartile range above the 75th percentile and below the 25th percentile. Jittered dots represent individual participants’ means. Solid gray lines connect two dots from the same participant. **(A)** Conflict effect in reaction times. **(B)** Conflict effect in mid-frontal (MF) theta power. **(C)** DCM amplitude estimates as obtained in Alameda et al. (2024) ^23^. **(D)** Linear model describing the effect of MF theta power on reaction times (RT). Both measures are conflict effect (I-C). **(E)** Linear model describing the effect of drift Diffusion Modelling for Conflict tasks (DMC) amplitude estimates on MF theta power. MF theta power represent the conflict effect (I-C).

The DMC estimate of automatic processes held a relationship with mean RT, suggesting that the effect of intensity level on MF theta power depends on task-irrelevant automatic processes. In fact, the model with the interaction between the predictors was the best fitting model, having the lowest AIC (23.173) and BIC (37.313). Specifically, increase in MF theta power was associated with an increase of interference effects from task-irrelevant features when fitting the data in the low-intensity condition (F(1,37) = 4.751, *p* = 0.036), but this relationship was lost when fitting the data in the high-intensity (F(1,37) = 0.237, *p* = 0.630) (Figure 3E). Further details about the methodology and the results of the exploratory analyses are available on the page of the preregistered report (https://osf.io/rdnmf).

### Multivariate spectral decoding

To overcome the shortcomings of univariate analysis and to capture more distribute networks, we turned to multivariate spectral decoding. Multivariate pattern analysis (MVPA) allow us to quantify conflict processing without ROI selection, capturing conflict-related changes harder to detect using conventional univariate analyses ^25^. MVPA on time-frequency data was able to decode congruency in the low-intensity state but not in the high-intensity state (Figure 4A). The significant cluster (p < 0.05, corrected for multiple comparison) appeared in the 219 - 609 ms time-window between 2 - 6 Hz with peak frequency at 6 Hz. The cluster was consistent with a congruency effect in the theta range. To further examine this effect, we run cluster-based permutation tests on the two conditions restricted to the cluster frequency range (2 - 6 Hz). In the low-intensity state only the cluster showed above chance classification between 266 - 594 ms (p < 0.05, corrected for multiple comparison), peaking at 359 ms (Figure 4B).

**Figure 4:**
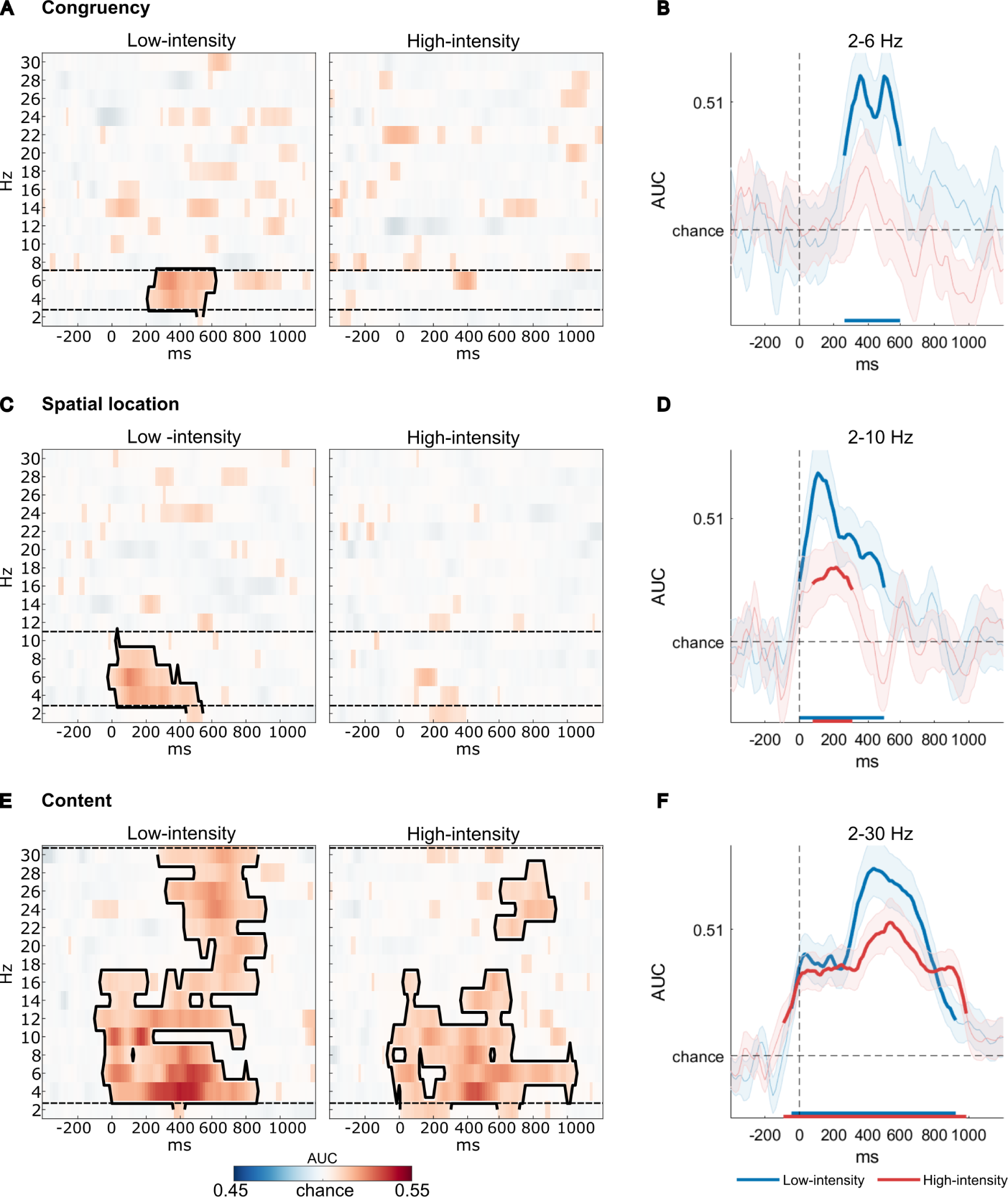
Multivariate spectral decoding results. Plotting of classifier accuracy against chance across the 2-30 Hz spectrum and all timepoints separately for low-intensity and high-intensity. Significant clusters (p < 0.05, corrected for multiple comparisons) are delimited by a solid black line. Dashed lines represent the minimum and maximum limits of significant clusters. The Area Under the Curve (AUC) within those limits in then averaged to depict the time course of classifier accuracy for low- and high-intensity (right column). Thicker lines represent statistical significance at p < 0.05. **(A)** Classifier accuracy for stimulus congruency. Information about congruency, and therefore the conflict effect, was only present at low-intensity in the 2-6 Hz band. **(B)** Time course of congruency decoding in the 2-6 Hz band in low- and high-intensity. **(C)** Classifier accuracy for stimulus physical location. Information about location was only present at low-intensity in the 2-10 Hz band. **(D)** Time course of spatial location decoding in the 2-6 Hz band in low- and high-intensity. Narrowing the cluster permutation analysis show that information about stimulus location is still preserved to some extent at high-intensity. **(E)** Classifier accuracy for stimulus content. Information about location was present in both conditions throughout the frequency spectrum. **(F)** Time course of content decoding throughout the whole spectrum (2-30 Hz) in low- and high-intensity.

Similarly, spatial location of the stimuli was decoded in time-frequency data at low-intensity (p < 0.05, corrected for multiple comparison). The cluster appeared in the -16 - 531 ms time-window between 2 - 10 Hz (peak frequency at 6 Hz), but not at high-intensity (Figure 4C). Cluster-based permutation testing restricted to the cluster frequency range (2 - 10 Hz) showed above chance classification in the low-intensity state between 0 - 500 ms (p < 0.05, corrected for multiple comparison), peaking at 109 ms. Notably, also high-intensity showed above chance classification between 78 - 313 ms peaking at 219 ms suggesting that some representation of physical features is preserved at high-intensity (Figure 4D). On the other hand, stimulus content was reliably decoded at both intensity conditions (Figure 4E). At low-intensity, content decoding was reliable between 2 - 30 Hz in the -94 - 906 ms time-window with peak frequency at 4 Hz (p < 0.05, corrected for multiple comparison). At high-intensity content decoding was reliable in two time-frequency regions. One between 2 - 16 Hz in the -63 - 1047 ms time-window with peak frequency at 4 Hz, the second between 22 - 28 Hz in the 578 - 922 ms time-window with peak frequency at 24 Hz. Cluster-based permutation testing averaging across the 2 - 30 Hz band showed a significant cluster at low-intensity -47 - 922 ms (p < 0.05, corrected for multiple comparison), peaking at 438 ms. At high-intensity, a cluster was significant between -94 - 984 ms, peaking at 531 ms (Figure 4F).

### Theta-band information sharing

To capture more drastic network reconfiguration during high-intensity, we performed weighted symbolic mutual information (wSMI). The time-window for wSMI analysis was 270-720 ms based on the significant interaction between intensity and congruency in MF theta (Figure 2A). We used a *τ* of 32ms to captures nonlinear information integration in the theta-band domain (4–9 Hz). The repeated measure ANOVA showed a main effect of congruency (F(1,38) = 7.831, *p* < 0.008, 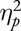 = 0.171, BF_10_ = 2.323). Neither the main effect of intensity (F(1,38) = 0.310, *p* = 0.581, 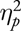 = 0.008, BF_10_ = 0.254) or the interaction were significant (F(1,38) = 0.011, *p* = 0.917, 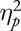 = 0.0002, BF_10_ = 0.232). Results are represented in Figure 5.

**Figure 5:**
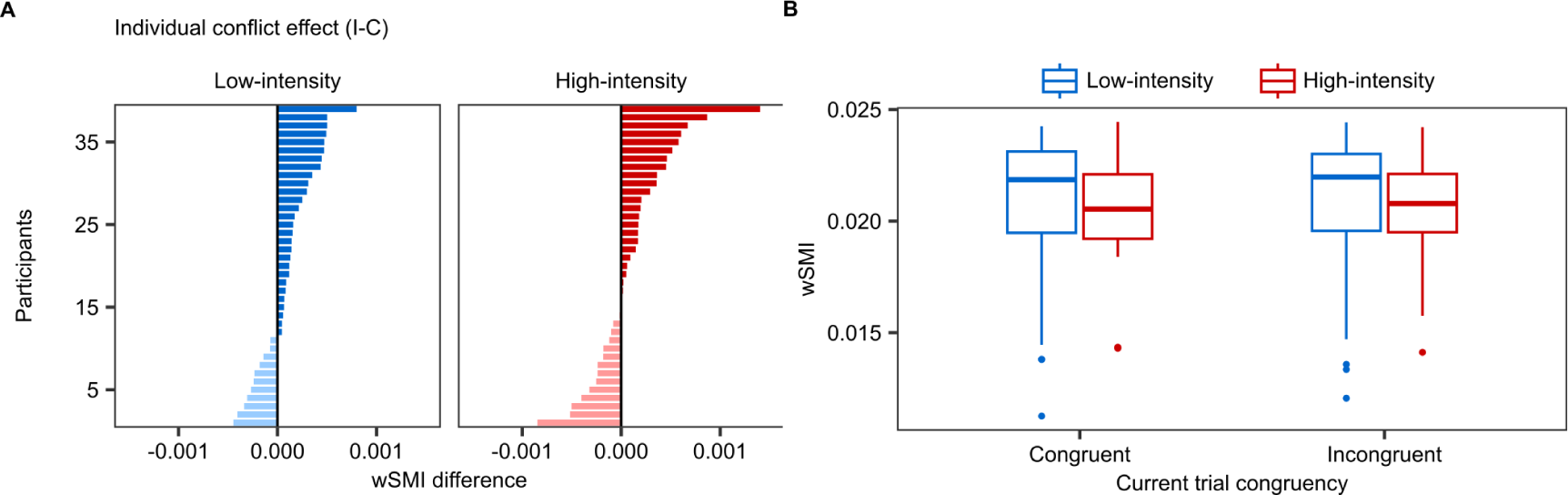
Theta-band information sharing results. **(A)** Individual weighted Symbolic Mutual Information (wSMI) conflict effects in the low- and high-intensity conditions. **(B)** Boxplots illustrating the effect of intensity level on the wSMI in response to congruent and incongruent trials. Boxplots show the median (horizontal line within each box), the interquartile range (IQR; the box), and 1.5 times the IQR (whiskers). Outliers are indicated as individual points beyond the whiskers. wSMI was calculated between each mid-frontal (MF) region of interest (ROI) electrode and every other electrode outside the ROI. We discarded from the analysis wSMI values within the ROI so that to evaluate only information integration between distant electrode pairs. The time window where we applied the wSMI was chosen based on the significant cluster representing the MF theta frequency interaction.

## DISCUSSION

The present study aimed at investigating behavioural and neural correlates of cognitive control during high physical and physiological activation induced by exercise. In line with our preregistered hypotheses, behavioural performance was preserved at high physiological activation. At the individual level, the effects of conflict and conflict adaptation were reliably detected. At the group level, the absence of interaction between exercise intensity and conflict effects showed that conflict processing was not hampered during high physiological activation. These results are consistent with previous literature showing that physical and physiological activation do not hinder performance in stimulus-response conflict tasks during acute exercise, even if at high intensity ^12,17,18^. Our results mirror those obtained in drowsy participants performing an auditory Simon task, in which the behavioural conflict effect was reliably detected even under reduced alertness ^10^. We did not, however, detect an overall slowing of RTs as a function of increased physiological activation. This confirms previous findings showing that cognitive control is very robust under high-intensity exercise ^17^. An alternative explanation is that measures of central tendency may provide only a gross representation of cognitive performance during acute exercise ^12,23^. In any case, consistently with the literature on both exercise and drowsiness, we show that even at high physiological activation individuals’ ability to resolve cognitive conflict remains effective.

As hypothesized, the robust behavioural conflict effect was not accompanied by an equally robust neural correlate. Indeed, high physical and physiological activation was associated with the degradation of neural signatures of conflict processing. The interaction between intensity and trial congruency in MF theta power suggests that conflict-induced power changes might be modulated by physiological activation. This effect was driven by the conflict effect being no longer reliable during high-intensity. Specifically, individual data (2C) showed that the conflict effect in MF theta was not only reduced in power but some participants even exhibited a reverse effect. MF theta power is considered one of the main neural indexes of cognitive control and a measure for processing demands across various cognitive interference tasks ^26–28^. Higher frontal theta power has been associated with preparation leading to better and faster conflict resolution ^29^.The increase in theta power in response to incongruent stimuli would reflect the additional recruitment of resources needed to resolve cognitive conflict ^30^. This additional recruitment was not detected at high-intensity in the present study. Or, at least, is was not detected by changes in theta power at MF locations. Alternatively, changes in MF theta power may reflect proactive control when initiating a motor response ^31–33^. In fact, MF theta has been shown to predict slower RTs in both conflict and non-conflict trials ^29^, possibly representing more deliberate control of motor movements. This suggests that increases in MF theta may not be conflict-specific but instead reflect a process more prevalent in, rather than exclusive to, conflict situations ^29,34^. Therefore, the absence of MF theta conflict effect may suggest that high-activation induces an overall increase in deliberate motor control regardless of conflict.

Exploratory analyses appear to further support that at high-intensity MF theta did not bear relationship with behavioural measures of cognitive control. In contrast, while at low-intensity MF theta power changes did not show a relationship with mean RT, it did with the DMC estimate of automatic processes, i.e., increase in MF theta power was associated with an increase of interference effects from task-irrelevant features under low-intensity condition, but this relationship was lost during high-intensity. This might suggest that the behavioural robustness of cognitive control is linked to decreased interference effect of task-irrelevant features ^23^ and that such link captured by theta power measured at MF locations is no longer reliable during high physiological activation. MF areas have been proposed to be the generator of conflict-modulated theta and changes in phase synchronization between MF, prefrontal and parietal regions within a frontoparietal network have been linked to cognitive control ^27,30,35^. MF regions would act as a central node that coordinates other regions and systems to monitor and adapt goal-directed behaviour essential for conflict processing ^36–40^. Focusing on the MF ROI, albeit justified by previous work, carries the drawback of unvariate analysis. This approach ignores the relevance of the multiple individual electrodes in differentiating between experimental conditions and overlooks the potential benefits of utilizing the covariance between electrodes. This oversimplifies complex interactions and undermines the role of neural networks involved in cognitive processes ^41,42^. The disappearance of the conflict effect in MF theta at high-intensity in conjunction with a preserved behavioural performance might reflect the rise of a more distributed network no longer detectable by a univariate approach. Alternatively, the absence of a conflict effect in the high-intensity condition may imply that physical activation causes a generalized increase in resource recruitment, potentially leaving no physiological capacity for additional conflict-induced resource allocation. This would be in line with both the observations that, in cognitively demanding tasks, higher demands in top-down control do not necessarily induce increased MF theta power ^31^, and that the performance of high-intensity physical exercise has been associated with an overall increase in brain activity ^16,43^. However, it appears that the theta frequency band is the only band of the spectrum that does not show a significant increase in power compared to a resting state ^16,43^. Therefore, it does not seem likely that physical exercise is causing a drastic increase in power in the theta band that would prevent the observation of the potential conflict effect in MF locations. Finally, the unreliability of the MF theta conflict effect in high-intensity might have also been influenced by noise. This is possible given the methodological challenges of a biking paradigm. For this reason, we conducted the experiment on a stationary bike with a fixed intensity and tested a sample of well-trained cyclists accustomed to maintaining a fixed posture for extended periods to minimize movement-related artifacts. Importantly, we did in fact replicate what was found in the drowsy state, where reduced physical and physiological activation was accompanied by the disappearance of the MF theta conflict effect ^10^. Following the same approach used to study cognitive control in drowsiness ^10^, we then implemented multivariate decoding which is fairly robust to noise as classifiers model the noise in the data by assigning low weight to non-informative artefacts ^44^. Importantly, by performing decoding within each intensity condition, noise level was equal for congruent and incongruent trials and for the training and testing phases.

Congruency decoding was consistent with the univariate MF theta. Here, decoding showed that information about stimulus congruency was reliably represented in neural data at low-intensity but not at high-intensity, similarly to what found in states of reduced (i.e., drowsiness) activation ^10^. The fact that decoding of congruency was preserved during low-intensity but not during high-intensity exercise further substantiate that changes in physiological activation levels are associated with changes in the neural mechanisms supervising the resolution of conflict. However, consistent with our hypotheses, even time-frequency multivariate analysis did not capture the neural signatures of conflict when the level of physiological activation increased. We hypothesized that the reconfiguration of these networks may be so pronounced to escape the sensitivity of multivariate decoding. We therefore implemented connectivity analysis based on information theory that capture changes in a wide network of brain regions rather than changes in local power ^10,45^. Specifically, we computed wSMI, that has been shown to reflect nonlinear brain inter-regional interactions ^46–48^. We observed a recovered conflict effect in wSMI at high-intensity which hints at the possibility that at high physiological activation the brain undergo the engagement of wider neuronal networks to resolve cognitive conflict. However, the findings from connectivity measures did not fully meet our hypotheses, as the wSMI analysis only showed a statistically significant main effect of congruency, suggesting that the effect was present both in the high- and low-intensity conditions. This leaves open the question of why a reliable congruency effect did not emerge in the univariate and multivariate EEG analyses during high-intensity. The analysis of the processing of the most relevant stimulus features in the present study may provide insights into this neural differentiation between the high- and low-intensity conditions.

At high-intensity, stimulus content was decoded above chance levels, showing that such information was retained. On the other hand, decoding of stimulus location was highly degraded, although when looking between 2-10Hz it appeared that some location information was retained. This finding was unexpected. A possible explanation may be that the functional organization of the auditory cortex is tonotopic rather than topographic ^49,50^, making the representation of spatial information less reliable in audition. In addition, the fact that spatial information was task-irrelevant may have contributed to the loss of its representation due to its limited importance for task performance. Hence, we consider the possibility that high physiological activation affected even perceptual processes such as the processing of spatial location, at least in the auditory modality.

With this study we propose that under heightened strained states reconfiguration of networks supervising cognitive control may occur. Whilst the interplay between physiological activation, cognition and its neural correlates have been traditionally studied in states of reduced activation ^7,8,10^, we provide a complementary view at the higher end of the physiological activation spectrum. Importantly, we focused on spontaneous fluctuations by inducing transitions towards high physiological activation in a natural way through exercise. High physiological activation through aerobic exercise is characterised by changes in the somatic and autonomic systems which are obtained by stressing the body with muscular, cardiovascular and respiratory demand. Similarly, somatic and autonomic changes happen when transitioning towards the lower end of the spectrum ^2^. By looking at natural physiological fluctuations (such as physical exertion and drowsiness), we suggest a framework for understanding the interplay between cognitive dynamics and physical activation. We propose that naturally occurring alterations in activation levels can be used to study the modulation of neural function and cognitive processing, going beyond traditional approaches of pharmacological interventions ^51^ or transient emotional state alterations ^52^. Traditionally, pharmacological and lesion-induced brain perturbations are considered strong causal methods in cognitive neuroscience. Nevertheless, physical/physiological activation, as an internally modulated change, can also serve as a powerful tool for studying cognition. Importantly, when natural fluctuations go undetected or are disregarded ^53^, they may potentially obscure or distort cognitive and neural markers of critical aspects of information processing ^7^. Utilizing physical activation and drowsiness as causal models to investigate the neural mechanisms of cognitive control may prove highly beneficial for exploring how cognition is hindered or remain resilient under naturally occurring physiological fluctuations. Specifically, here we suggest that when individuals are asked to perform a cognitive task, both states may act as stressors on the system, necessitating a greater recruitment of resources to efficiently perform the task compared to optimal states (such as awake and resting). Importantly, the two physiological states do not necessarily represent opposite ends of the same linear spectrum and the changes that occur under high physical demand are not necessarily linearly related to what occurs in states of reduced activation. Rather, they may share the role of stressors on the system, engaging similar compensatory mechanisms to overcome the challenges that physiological fluctuations pose to cognitive performance. In sum, we suggest that these states might share some common processes in the way that cognition and behaviour are supervised by the brain. Clarifying the nature of the commonalities at the opposite ends of the physiological activation spectrum will be an exciting task for future research.

## METHODS

### Participants

Thirty-nine healthy subjects (2 females; age-range= 18-50 years; mean age = 30.55 years; SD age = 10.65; 4 left-handed) were included for analyses. Initially, forty participants were recruited, but one participant was excluded due to not meeting the minimum average heart rate criterion at high intensity. This criterion was set at 75% of the individual’s maximum heart rate. Participants were recruited by word of mouth and written advertisement amongst cycling and triathlon sport clubs, and received 30 euros for their participation. Only expert cyclists were included in the study. They were required to have a minimum of three years experience and to train at least three times a week. All participants had normal or corrected-to-normal vision, normal hearing, no history of head injury or physical and mental illness. This study was approved by the ethical committee of the University of Granada (978/CEIH/2019). Written informed consent was obtained from all participants during the first experimental session, after explanation of the experimental protocol. All ethical regulations relevant to human research participants were followed.

### Study design

To test the effect of physiological activation on the resolution of cognitive conflict, we adopted a 2 × 2 factorial design with exercise intensity level (low/high) and the congruency of the current trial (congruent/incongruent) as factors. We also investigated the moderating effect of previous trial congruency (conflict adaptation). We adopted a 2 × 2 × 2 design with intensity, current trial congruency, and previous trial congruency (pre-congruent/pre-incongruent) as factors. Note that for aiding interpretation, the factor congruency was in fact reduced to a measure of conguency effect by subtracting the signal elicited by congruent trials from the signal elicited by incongruent trials. All factors of both study designs were within-subjects. Analyses on conflict adaptation were run on behavioural data only as an additional sanity check.

### Experimental task

Participants performed an auditory version of the Simon task ^22^. Recordings of the spoken words “left” and “right” (”izquierda” and “derecha” in Spanish) were played monoaurally through in-ear earphones. In congruent trials (50% of the trials), the stimulus was presented at the physical location corresponding to the content of the word (i.e. “left” played to the left eat, “right” played to the right ear). In incongruent trials, physical location and content did not correspond (i.e. “left” played to the right ear, “right” played to the left ear). Participants were instructed to respond according to the content of the word, while ignoring its physical location. Response was given with the left and right thumbs by pressing one of two buttons that were attached to the bike handle (Figure 6A). Speed and accuracy were stressed. Interstimulus interval (ITI) between participant’s response and the next stimulus onset was a random number between 1500 and 2000 ms. If the participant did not respond, the ITI began 1900 ms after stimulus onset (Figure 6B). Practice and testing blocks were fixed in time duration, therefore the total number of trials depended on the participant’s speed. The practice block lasted 30 s and on average participants responded to 12 trials (SD = 1). The testing block lasted 30 min without breaks and on average participants responded to 766 trials (SD = 41). Each participant completed an equal number of trials requiring responses with the left and right hand. The trials were equally divided among the four conditions: congruent-right, congruent-left, incongruent-right, and incongruent-left, each comprising one quarter of the total trials. Although the total number of trials varied due to the participants’ response speeds, there were no significant differences in the number of trials requiring either hand. The same presentation script was used for both the low and high-intensity sessions to ensure consistency. A beep sound was played every 10 min to inform the participant of the elapsed time.

**Figure 6:**
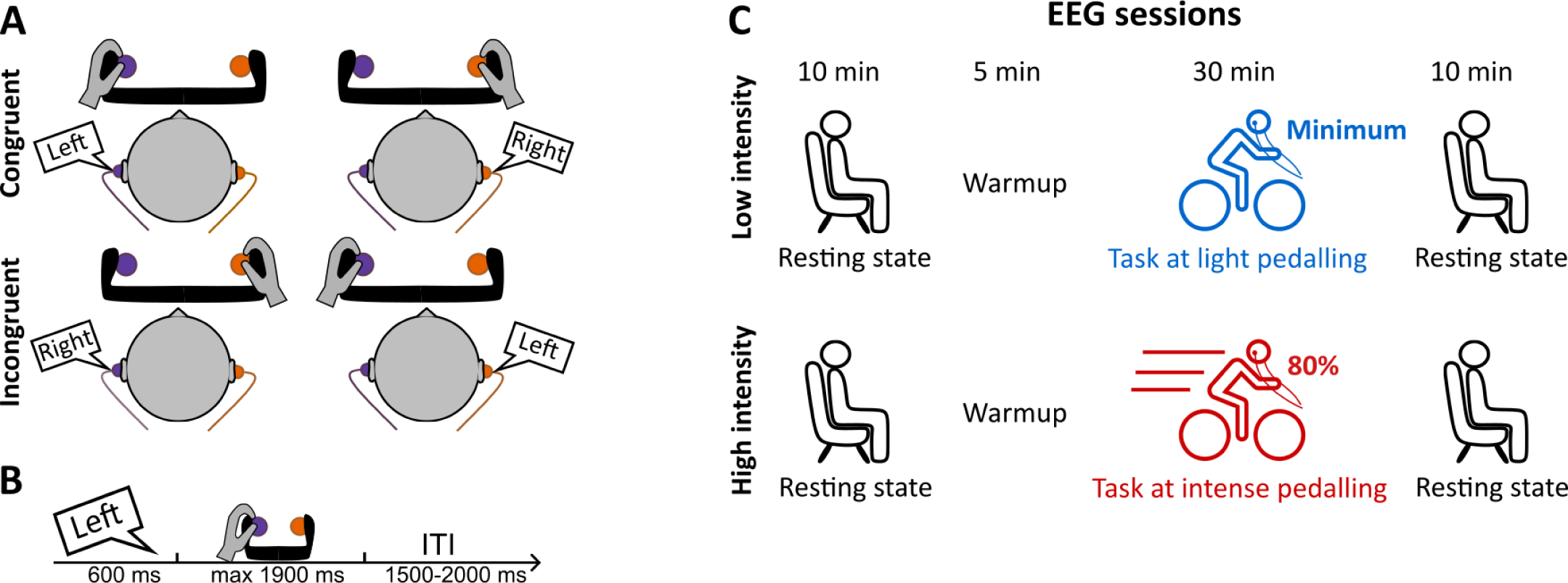
Task and procedure. **(A)** Participants were presented with the words “left” and “right” in their left and right ear. The task required to respond to the content of the word by pressing a button with their right or left thumb while ignoring the stimulus spatial location. Stimulus content and spatial location could be either congruent or incongruent. **(B)** Example of trial structure. **(C)** The session started with a seated 10 min resting state. After that, participants mounted on the bike and warmed up for 5 min, during which a 30 s practice block was presented. Then, the experimental task was presented for 30 min during which participants had to pedal without stopping. In the low-intensity condition, the resistance of the bike was set at 10% of the maximum W that the participant had moved in the effort test. In the high-intensity condition, the resistance was set at 80%. The session ended with a seated 10 min resting state. Note that in this paper resting state data are not analysed.

### Procedure

Testing took place over three sessions: the first was always a maximum effort test, the second and third were EEG experimental sessions. Upon arrival for the effort test, participants were informed on the overall experiment including that one of the subsequent EEG sessions would have been at high-intensity and one at low-intensity. Then, they gave written consent. The order of the EEG sessions was counterbalanced across participants and it was not communicated until the first EEG session. Participants were instructed not to perform intense exercise during the 48 hours preceding any of the three sessions. The maximum effort test took place about one week before the first EEG session, while the EEG sessions were 48 to 72 hours apart to avoid fatigue effects. To accommodate for participants life commitments, four participants had more than 72 hours in between EEG sessions, and three only had 24 hours. In the case of the three latter, the low-intensity session was done first. Finally, to control for circadian rhythms, each participant performed both EEG sessions at the same time of day. However, for practical purposes, different participants were scheduled at various times. Testing sessions commenced approximately between 9:00 and 11:00 (10 subjects), 12:00 and 13:00 (5 subjects), and 16:00 and 18:00 (14 subjects).

### Maximum effort test

The maximum effort test consisted of an incremental cycle-ergometer test (ergoline GmbH, Germany) to confirm participants’ fitness and individually adjust the exercise intensity in the following EEG sessions. Participants were fitted with a wireless 12-lead PC ECG (AMEDTEC Medizintechnik Aue GmbH, Germany) to record HR and with a gas analyser mask (Geratherm Respiratory GmbH, Germany) to measure VO2. Participants were informed about the protocol and instructed to keep pedalling at above 60 rpm until they reached full exhaustion and could no longer keep going. The protocol started with 5 min warmup at 120 watts (W) immediately followed by the incremental test of 20 W every 3 min. For each individual participant, the power output in W corresponding to when they reached their maximum oxygen consumption was used as marker to calculate the intensity of the following EEG sessions.

### EEG sessions

Upon arrival, participants were seated in a comfortable chair in an electrically shielded room, and were fitted with a 64-channels EEG cap (EASYCAP GmbH, Germany) and three ECG electrodes (Ambu A/S, Denmark). During electrodes placement, participants received task instructions in written form. The session started with 10 min resting state. Note that resting state data were collected for a different study and therefore are not included in the manuscript. Then, participants mounted on a stationary bike (SRM International SRM GmbH, Germany) and fitted with a heart rate monitor SRM HR (SRM International SRM GmbH, Germany) and in-ear earphones (Hyperx, HP Inc., USA). Participants were reminded to respond as accurately and fast as possible and to keep a cadence of 60 rpm or above during testing. Participants were first given 5 min to warm up on the bike. In the warmup before the low-intensity session, the resistance of the bike was set at the minimum load possible while preventing the wheel from spin freely. For the high-intensity session, the warmup was incremental, with resistance increases of 20 W every minute with the last step being the 80% of the W at VO2_max_. During the warmup of the first session, the practice block was presented. Then, the experimental test began during which participants performed the auditory Simon task while pedalling (Figure 6C). In the low-intensity session, the resistance of the bike was set at the minimum load and in the high-intensity session at 80%. Given the importance of finishing the session and the heavy physiological demand of the high-intensity session, W were adjusted manually if the participants’ cadence dropped below 60rpm or if their HR exceeded the 90% of their HR max as measured during the effort test. The resistance of the stationary bike was controlled with Gobat-Software (Magnetic Days O.R.F. s.r.l., Italy). W output and HR were recorded with SRM powercontrol 8 (SRM International SRM GmbH, Germany). The session ended with 10 min resting state not included in the current manuscript. Stimuli were presented using PsychToolbox ^54,55^ software on a Windows computer.

### Behavioural data analysis

The first trial of each session, incorrect responses, omissions and responses faster than 200 ms were removed. Analyses were conducted on both RT and ACC. To test the conflict effect, we run 2 (intensity level) × 2 (congruency) repeated measure ANOVAs. Post-hoc planned comparisons corrected for multiple comparisons were conducted comparing congruent vs incongruent in low-intensity and high-intensity. To test the conflict adaptation effect, we run 2 (intensity level) × 2 (pre-congruency) repeated measure ANOVAs with the conflict effect (incongruent minus congruent) as dependent variable. Post-hoc planned comparisons corrected for multiple comparisons compared pre-congruent vs pre-incongruent in low-intensity and high-intensity. In addition to the frequentist approach, we also performed the correspective bayesian ANOVAs and planned comparisons.

### EEG recording and preprocessing

64-channel EEG data sampled at 1000 Hz and referenced to FCz according to the extended 10-20 system were collected using the actiCHamp amplifier and Brain Vision software (Brain Products GmbH, Munich, Germany). The ground electrode was placed at the centre of the forehead according to the actiCAP configuration. Electrode impedances were kept below 10 kΩ. EEG preprocessing was conducted using custom MATLAB scripts, the EEGLAB ^56^, and FieldTrip ^57^ toolboxes. Continuous data were dowsampled to 500Hz, notch-filtered and band-pass filtered 0.5 Hz-90 Hz. We filtered the data using Hamming windowed FIR filters as implemented in EEGLAB ^56^. For high-pass, the filter order and transition band width was calculated as the 25% of the lower passband edge, but not lower than 2 Hz. For low-pass it was determined by the distance from the passband edge to the critical frequency (Nyquist). The cut-off frequency was defined at -6 dB. Bad channels were detected using joint probability with a threshold set at *±* 3.5 SD. For each individual session, Infomax-based Independent Component Analysis (ICA) was then run on good channels and continuous data. The infomax algorithm maximizes the entropy of the output of a neural network, which leads to the maximization of the mutual information between the input and the output ^58^. This process results in the extraction of statistically independent sources from a set of mixed signals and allows to separate artifacts (such as eye blinks or muscle activity) from neural signals. The EEGLab runica algorithm was applied with default parameters. We used the ICLabel EEGLAB plugin to identify ICA components that were classified as having above 80% probability to be generated by eye-movements, muscle, or heart activity. To avoid the effects that filtering can have on MVPA classification ^59^, those components were then rejected from unfiltered data downsampled at 500Hz. The signal was then rereferenced to the average and bad channels were interpolated. The continuous recording was epoched between -1500 and +2000 ms relative to stimulus onset. The epochs corresponding to those trials that had been removed from behavioural data were rejected. To further clean from eye movements and muscle activity, we rejected abnormal spectra epochs whose spectral power deviated from the mean by *±*50 dB in the 0–2 Hz frequency band and by +25 or −100 dB in the 20–40 Hz frequency band. After trial rejection, an average of 710.2 (SD = 89.199) trials per participant were retained in the high-intensity condition and an average of 713.05 (SD = 82.817) per participant were retained in the low-intensity condition.

### EEG time-frequency analysis

Epochs were grouped based on current trial congruency. Time-frequency decomposition was run using FieldTrip ^57^. Morlet wavelets of 3 cycles (the cycles number remained constant throughout frequencies) were adopted to conduct trial by trial time-frequency decomposition. The signal was decomposed between 2 and 30 Hz in linear steps of 2 Hz. The wavelet slid from -400 to 1250 ms (relative to stimulus onset) in steps of 10 ms. Then, the epochs were transformed into decibels (dB) and normalized to a baseline of -400 to -100 ms relative to stimulus onset. For each participant, epochs were averaged according to the two trial types. Then, the signal corresponding to the MF ROI was selected and averaged across electrodes (C1, Cz, C2, FC1, FCz, FC2, F1, Fz, F2. See Figure 2A). We obtained the time-frequency conflict effect at the participant level by subtracting the signal to congruent trials from the signal to incongruent trials. This allowed us to use cluster-based permutation testing ^57^ to directly test the interaction between congruency and intensity level by comparing the conflict effect between high- and low-intensity. For the cluster-based permutation tests, the Monte Carlo method was implemented with 1000 iterations and the alpha threshold at the cluster level was set at 0.05. The test were run on all frequencies and timepoints between -200 ms and 1200 ms. To better interpret the effect of the interaction, we conducted dependent samples t-statistic with alpha at 0.05. Finally, during the review process, the need arose for a deeper analysis of the interaction to understand the driving factors behind the observed effect. We therefore obtained the average power within the ROI in each condition and run 2 (intensity level) × 2 (congruency) repeated measure ANOVAs. Subsequently, we performed Bonferroni-corrected pairwise comparisons between low-intensity and high-intensity in the congruent and incongruent conditions. It is important to note that these additional analyses use power from an ROI that inherently reflects a significant interaction between intensity and congruency factors. Consequently, we caution that this constitutes a circular analysis ^60^, and that these analyses have been reported mainly to facilitate the interpretation of the cluster permutation tests.

### EEG multivariate spectral decoding

Multivariate spectral decoding was applied on the time–frequency data. Multivariate analysis increase sensitivity and allows to query how stimulus features are processed at different levels of intensity. To do so, we used the ADAM toolbox ^25^ which has been developed to specifically implement MVPA on EEG data. We input the raw data which were first down-sampled to 64 Hz and then converted to time-frequency directly by the ADAM toolbox using default options. Continuous data were epoched between -1500 ms to 2000 ms and in the frequency domain were decomposed between 2 Hz and 30 Hz in steps of 2 Hz. To avoid biased decoding, we balanced the number of trials between conditions per state of intensity. We performed three separate decoding analyses taking current trial congruency (congruent/incongruent), content (”right”/”left”), and physical location (right/left) as classes. 10-fold cross-validation was performed applying a backward decoding algorithm. Linear discriminant analysis (LDA) was used to discriminate between stimulus classes, then classification accuracy was computed as the area under the curve (AUC). Finally, AUC scores were tested per time point and frequency bin with double-sided t-tests against a 50% chance level across participants. To control for multiple comparisons, t-tests were conducted with cluster-based permutations with alpha 0.05 and 1000 permutations. Note that content and physical location decoding had not been preregistered. However, they follow the methodology of ^10^

### weighted Symbolic Mutual Information

We quantified long-range information sharing between electrodes by means of wSMI ^47,48,61^. The wSMI has the advantage of making a rapid and robust estimation of the entropy of the signals, detecting nonlinear coupling while discarding the spurious correlations between signals arising from common sources. For each trial, the EEG signal was transformed into a sequence of discrete symbols. Such transformation depends on the length of the symbols k and their temporal separation *τ*. We chose a kernel size (k) of 3, *τ* values of 32 ms to obtain sensitivity to frequencies spanning the theta band (4–9 Hz). For each pair of transformed signal, the wSMI was estimated by calculating the joint probability of each pair of symbols. The joint probability matrix was multiplied by binary weights to reduce spurious correlations between signals. For pairs of identical symbols that could be elicited by a common source, and for pairs of opposite symbols that could reflect the two sides of a single electric dipole, the weights were set to zero. wSMI was calculated between each MF ROI electrode and every other electrode outside the ROI. We discarded from the analysis the wSMI values within the ROI so that to evaluate only information integration between distant electrode pairs. The time-window where we applied the wSMI was chosen based on the significant cluster representing the MF theta frequency interaction.

### Statistics and Reproducibility

Statistical analyses were run with MATLAB 2019b and R version 4.2.1. Bayes factors are reported as the degree of evidence for the alternative over the null hypothesis (BF_10_). Bayes factors were calculated with JASP ^62^ with the null model as denominator and default settings. For the main effects of the F-statistics the null model included subject and random slopes, while for the interaction effects the null model included subject, random slopes, and the main factors intensity and congruency. Hypotheses and analyses plan had been preregistered prior to data collection ^21^. Please note that we expected the behavioural conflict adaptation effect results to follow the same pattern of those of the conflict effect, consistently with previous literature ^10^. The fact that the preregistered hypothesis on the behavioural conflict adaptation effect says otherwise, is a genuine mistake made at the time of the drafting of the preregistered report.

### Exploratory analyses: Relationship between MF theta power and behavioural measures

We conducted exploratory analyses to investigate the relationship between behavioural measures and MF theta power changes induced by arousal modulation. To do so, the MF theta power conflict effect was extracted separately for the high and low arousal condition. We subtracted the time-frequency decomposition from congruent trials from that from incongruent trials. Then we averaged the power within the significant cluster region that had been observed in the previous cluster-based permutation testing.

First we investigated whether MF power changes predicted changes in RT by taking into account individual variability. We hierarchically compared three nested models which all had the RT conflict effect as dependent variable and participants as random effect: model A0 was the intercept, model A1 included arousal level as predictor. Model A2 included arousal level and MF theta power as predictors. Model 3 included an interaction term between arousal level and MF theta power. The interaction term allow for potential variation in the effect of MF theta power on RT across participants.

Furthermore, we investigated whether MF theta power changes held a relationship with the interference of task-irrelevant automatic processes. In a study conducted on the same dataset as the one presented in the current paper ^23^, behavioural dynamics were modelled with Drift Diffusion Modelling for conflict tasks (DMC) ^24^. DMC assume that, when an individual faces a two-alternative choice, the amount of evidence for one answer over the other accumulates gradually over time until it reaches a decision threshold, which triggers the motor response ^63^. DMC allow to estimate the influence of both automatic (task-irrelevant) and control (task-relevant) processes on decision time. Alameda et al. (2024) ^23^ observed that the estimate representing automatic processes (the amplitude to peak of automatic processing pulse function) was reduced in high-arousal compared to low-arousal, suggesting that the amount of interference from non-relevant information would be slightly reduced during the high-arousal state. We therefore explored the possibility that MF power changes are predicted by latent psychological processes underlying decision making. In particular, that it would be predicted by state-dependent variations in the processing of task-irrelevant information. To do so, we estimated for each participant the amplitude to peak of automatic processing pulse function with DMCfun package in R ^64^ (for more details on the methodology, see the original article ^23^). Then, we fitted linear mixed effect models to investigate the effects of arousal lever and the DMC amplitude estimate on theta power, while accounting for individual variability (random effect). Specifically, we hierarchically compared three nested models, all with participant as random intercept: model B0 was the intercept, model B1 included DMC amplitude as predictor. The model B2 included DMC amplitude and arousal level as predictors. This model assumes that the relationship between power and the DMC amplitude estimate is linear and that the effect of DMC amplitude on power is constant across all participants. Model B3 included an interaction term between DMC amplitude and arousal level. The interaction term allow for potential variation in the effect of amplitude on power across participants.

We compared the fit of the models with likelihood ratio test, Akaike Information Criterion (AIC) and Bayesian Information Criterion (BIC) to determine which one best explained the data. Note that these analyses had not been preregistered and were planned upon observation of the current data and the results of Alameda et al (2024) ^23^. They are therefore exploratory and will be interpreted as such.

## DATA AVAILABILITY

The data generated and analysed during this study are available in BIDS format at the OSF preregistration page https://osf.io/rdnmf which will redirect to the Zenodo repository.

## CODE AVAILABILITY

The code used to analyse the data in this study are available at the OSF preregistration page https://osf.io/rdnmf which will redirect to the Zenodo repository. The code was written with MATLAB 2019b and R version 4.2.1. The toolboxes EEGLAB version 13_5_4b, FieldTrip version 20181231, and ADAM version 1.14-beta were used for EEG preprocessing.

## AUTHOR CONTRIBUTIONS

**Chiara Avancini**: Conceptualization, Methodology, Software, Data curation, Formal Analysis, Visualization, Writing, Reviewing and Editing. **Luis F. Ciria** Conceptualization, Methodology, Software and Reviewing. **Clara Alameda**: Data curation, Reviewing and Editing. **Ana F. Palenciano**: Methodology. **Andres Canales-Johnson**: Conceptualization, Methodology, Reviewing and Editing. **Tristan A. Bekinschtein**: Conceptualization, Methodology, Reviewing and Editing. **Daniel Sanabria**: Conceptualization, Methodology, Reviewing, Editing, and Funding Acquisition.

## ACKNOWLEDGEMENTS

This study was supported by a research project grant from the Spanish Ministry of Science and Innovation to Daniel Sanabria (PID2019-105635GB-I00); a postdoctoral fellowship by the Spanish Ministry for Science and Innovation awarded to Chiara Avancini (FJC2020-046310-I); a postdoctoral fellowship from the Regional Government of Andalusia awarded to Luis F. Ciria (DOC_00225); a predoctoral fellowship by the Spanish Ministry of Universities awarded to Clara Alameda (FPU21*/*00388). Andres Canales-Johnson is supported by an ANID/FONDECYT Regular (1240899) research grant. We thank Juan José Pérez Díaz for his assistance with the maximum effort tests.

